## Supplementary-Material_Lenschow_Mendes for "A galanin-positive population of lumbar spinal cord neurons modulates sexual arousal and copulatory behavior"

### Supplementary Materials

### Supplemental Figures

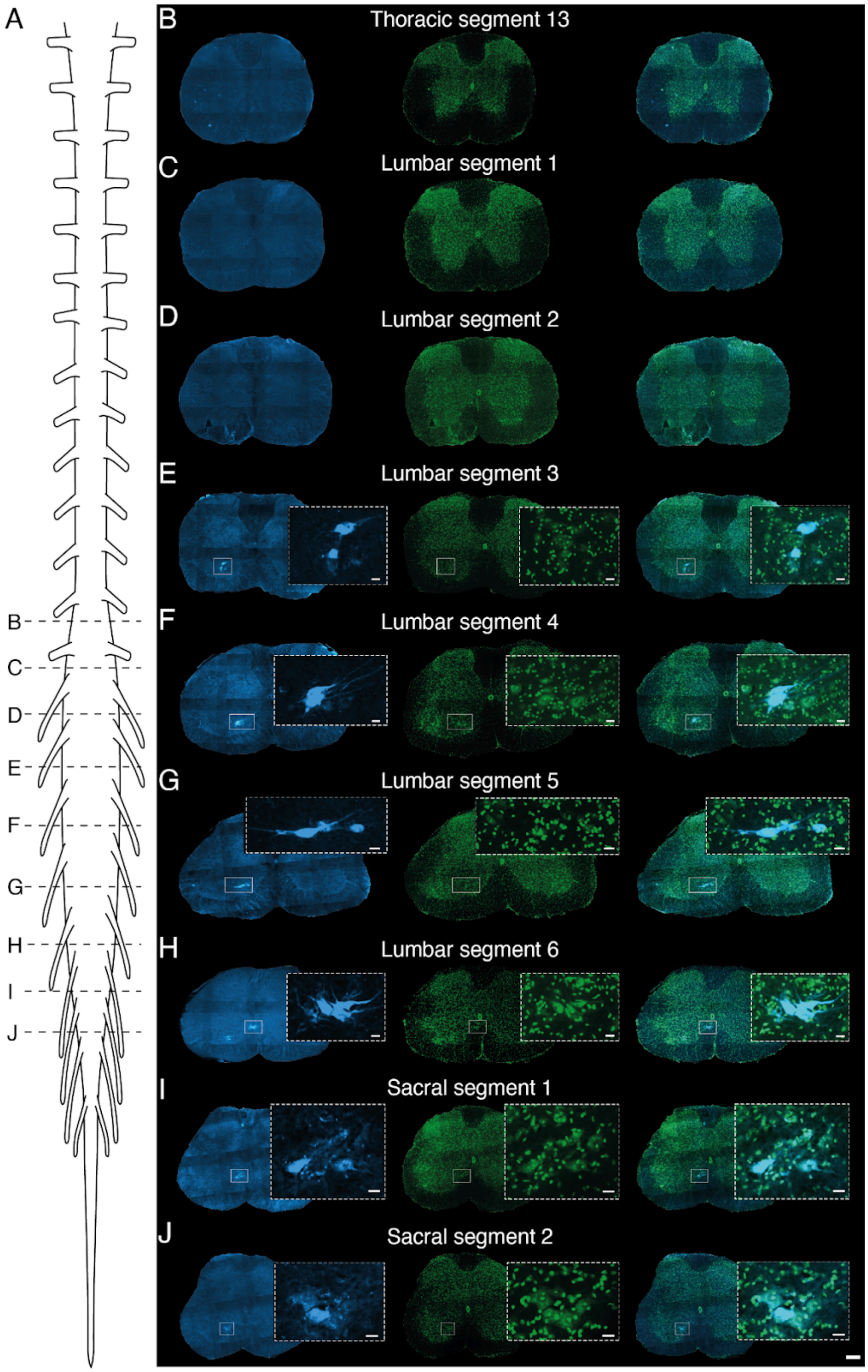

**Supplemental Figure 1: Anatomical distribution of the bulbospongiosus muscle motor neurons (BSM-MNs).**

(A) Spinal cord scheme illustrating segments that are shown in B-J.

(B - J) Left panels: Blue channel revealing FG labeled BSM-MNs (E -J). Middle panel: Green channel illustrating the Nissl stain in order to identify the spinal cord segments. Right panel: Overlap of both channels. Scale bar 200um. Inset scare bar: 50um.

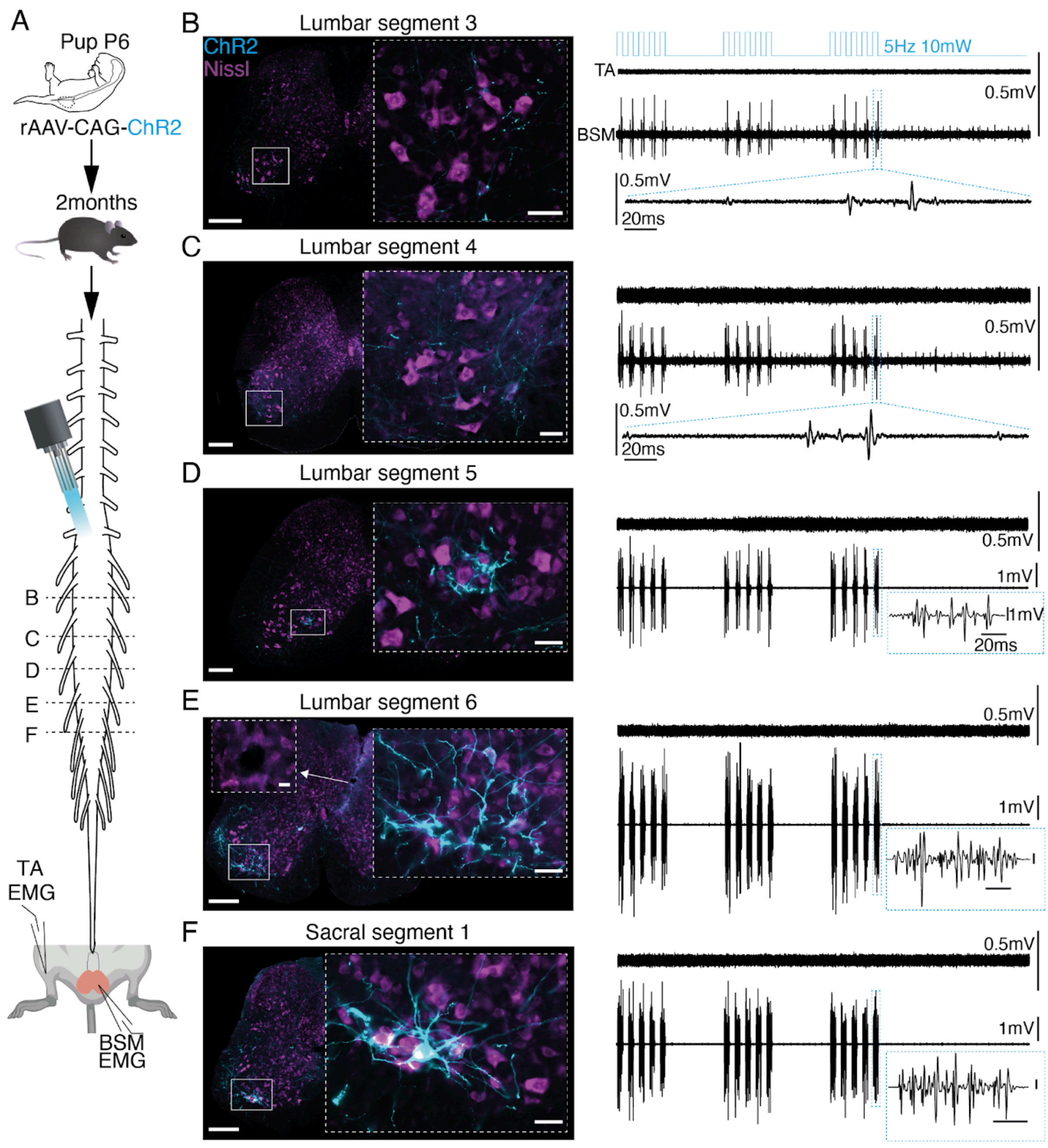


**Supplemental Figure 2: Optogenetic stimulation of the BSM-MNs along the rostrocaudal axis of the spinal cord.**

(A) Experimental setup. Left panel: BL6 mice pups (aged postnatal day 3-6) received an injection of a rAAV-CAG-ChR2 on the BSM. Pups were raised until 2-3 months of age, to perform optogenetic stimulation on top of the spinal cord (rostral to caudal) while in parallel monitoring EMG activity in the BSM.

(B) Leftpanel: Representative spinal cord segment (lumbar segment 3, L3) of a 2 month old mouse that was infected at the age of P4-P6 with a rAAV-CAG-ChR2 (blue) in the BSM. Identification of spinal cord segments based on the atlas was achieved via a Nissl stain (purple). Right panel: corresponding EMG recordings (upper trace: TA in the leg; lower trace: BSM) observed while shining blue light on top of the spinal cord segment shown in the left panel. EMG recordings were aligned to histology by placing an electrolytic lesion at the spot with the largest BSM response recorded (see panel E). Scale bar 200um. Scale bar inset 20um.

(C-F) Same as B for the remaining spinal cord segments depicted in A (see dotted lines, from L3-S1). Note that the biggest responses in the BSM EMG were triggered at the location where most BSM-MNs were infected (L6, S1).

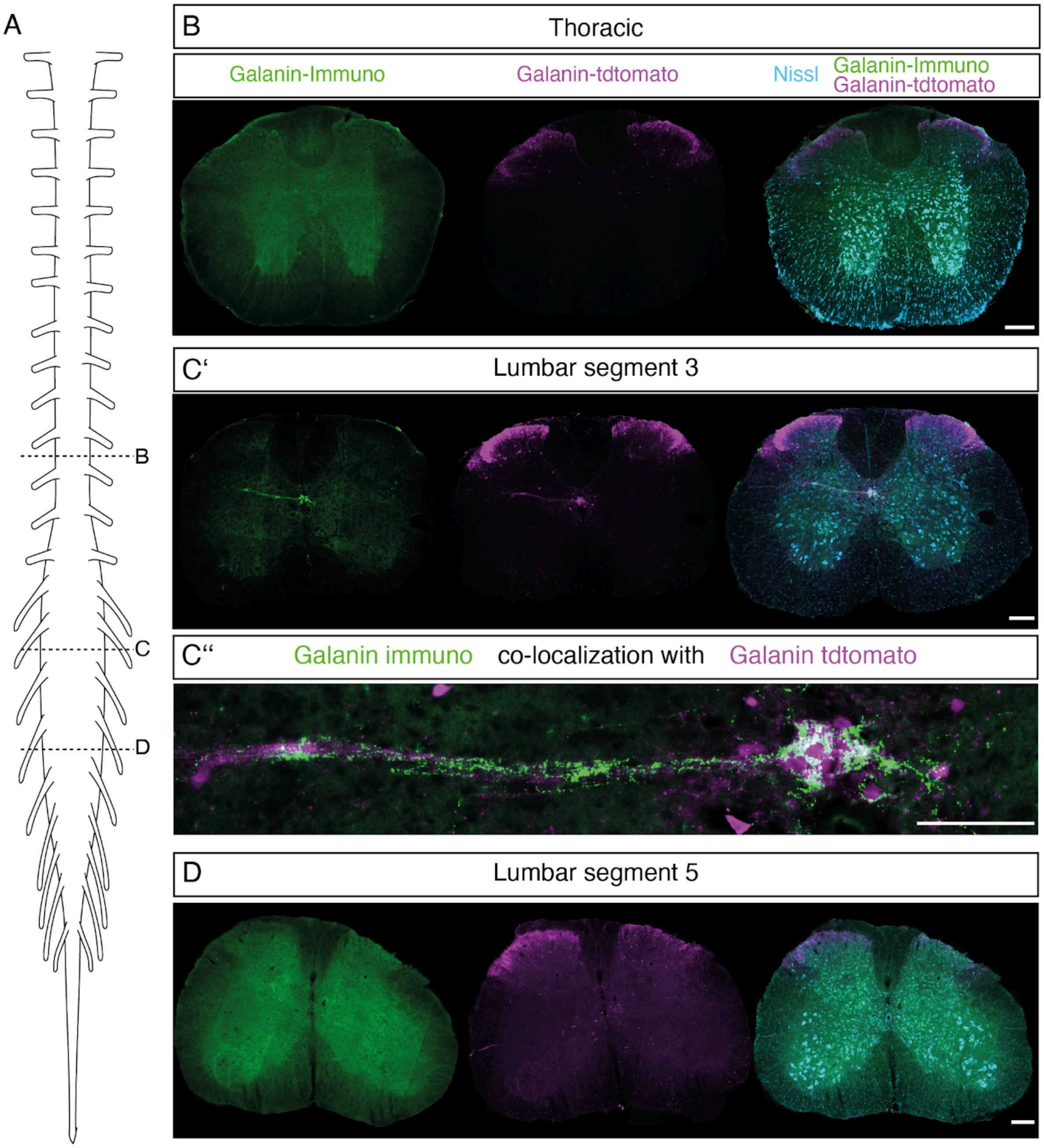


**Supplemental Figure 3: The Gal+ population is mostly present at the L2/3 spinal segments.**

(A) Scheme of spinal cord.

(B) Example staining, in the thoracic part depicted in A, for Galanin (left, green) in a Gal-cre x TdTomato animal (purple, middle). The Nissl stain (cyan) served as an identification parameter for the spinal cord segments. Note that there is no Galanin signal around the central canal. Scale bar 200um.

(C’) Same as B but for the L3 segment. Note the presence of a prominent cluster of Galanin expressing cells (shown with the immunostaining, green, and the tdTomato signal, purple, of the Galanin-reporter line) around the central canal.

(C”) Overlay of the Galanin immunohistochemical signal (green) with the tdTomato signal in the Gal-cre x TdTomato cross reveals an overlap of both signals (see zoom in). Scale bar 100um.

(D) Same as B but for the L5 spinal segment. Note again the absence of Galanin positive cells around the central canal.

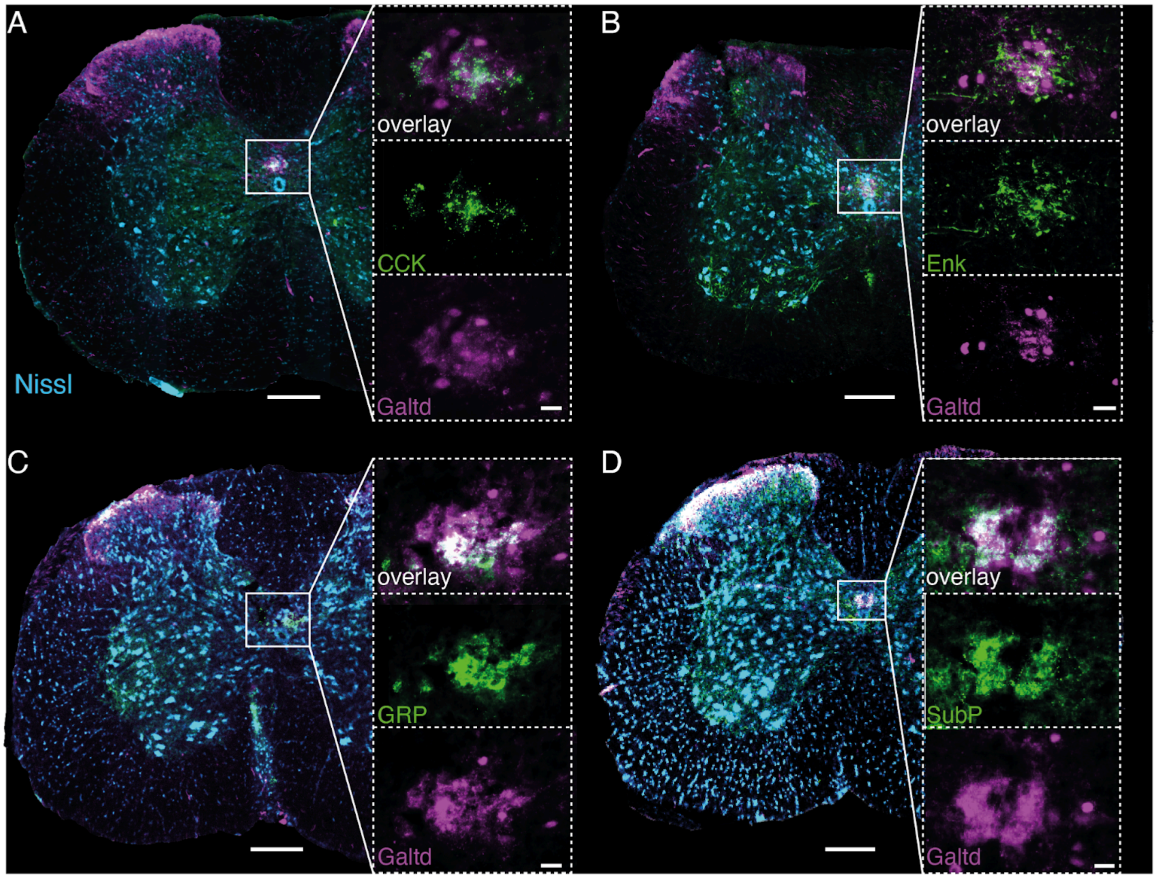


**Supplemental Figure 4: Immunohistochemical characterization of the lumbar Gal+ population.**

(A) Immunohistochemical staining for Cholecystokinin (CCK; green) in the L2/3 spinal segments obtained from a Gal-cre x dTomato (Gal-td, pink) male mouse. A Nissl stain (cyan) was performed to identify the spinal cord segment. Scale bar spinal segment 200um. Scale bar insets 20um.

(B) Same as A but immunohistochemical processing was done for Enkephalin (Enk).

(C) Same as A but a post-hoc staining for Gastrin-releasing peptide (GRP) was performed.

(D) Same as A but immunohistochemical staining against Substance P (SubP) was performed.

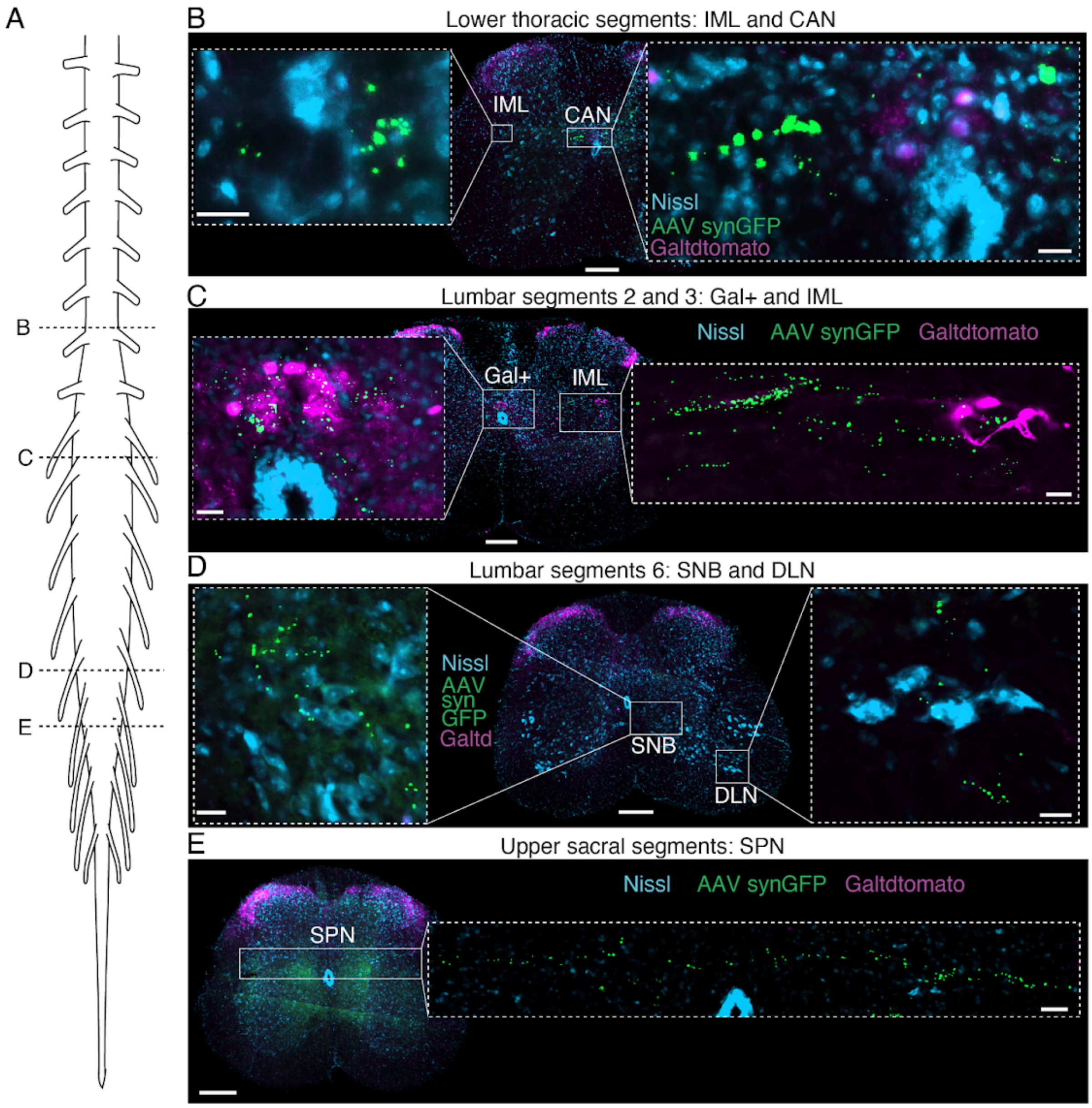


**Supplemental Figure 5: Spinal projection targets of the Gal+ cells revealed with a marker for postsynaptic boutons (synaptophysin delivered through an AAV-flex-CAG-SynGFP) .**

(A) Scheme of spinal cord.

(B) GFP-labeled synaptic terminals of Gal+ cells were found at the location of the central autonomic nucleus (CAN) and intermediolateral nucleus (IML) at lower thoracic segments. Both nuclei are known to contain sympathetic preganglionic cells. Scale bar 200um. Scale bar insets 20um.

(C) Prominent GFP-labeled boutons were found at the location of Gal+ cells and at the IML of the L2/3 spinal segments. Scale bar 200um. Scale bar inset 50um.

(D) At the L6 segment, GFP-positive synaptic terminals were dominantly found at the spinal nucleus of bulbocavernosus (SNB) that contains the BSM-MNs and at the dorsolateral nucleus (DLN) which contains the MNs controlling the ischiocavernosus muscle.

(E) In upper sacral segments GFP-positive terminals were diffusely found in regions containing parasympathetic preganglionic cells (SPN, sacral parasympathetic nucleus).

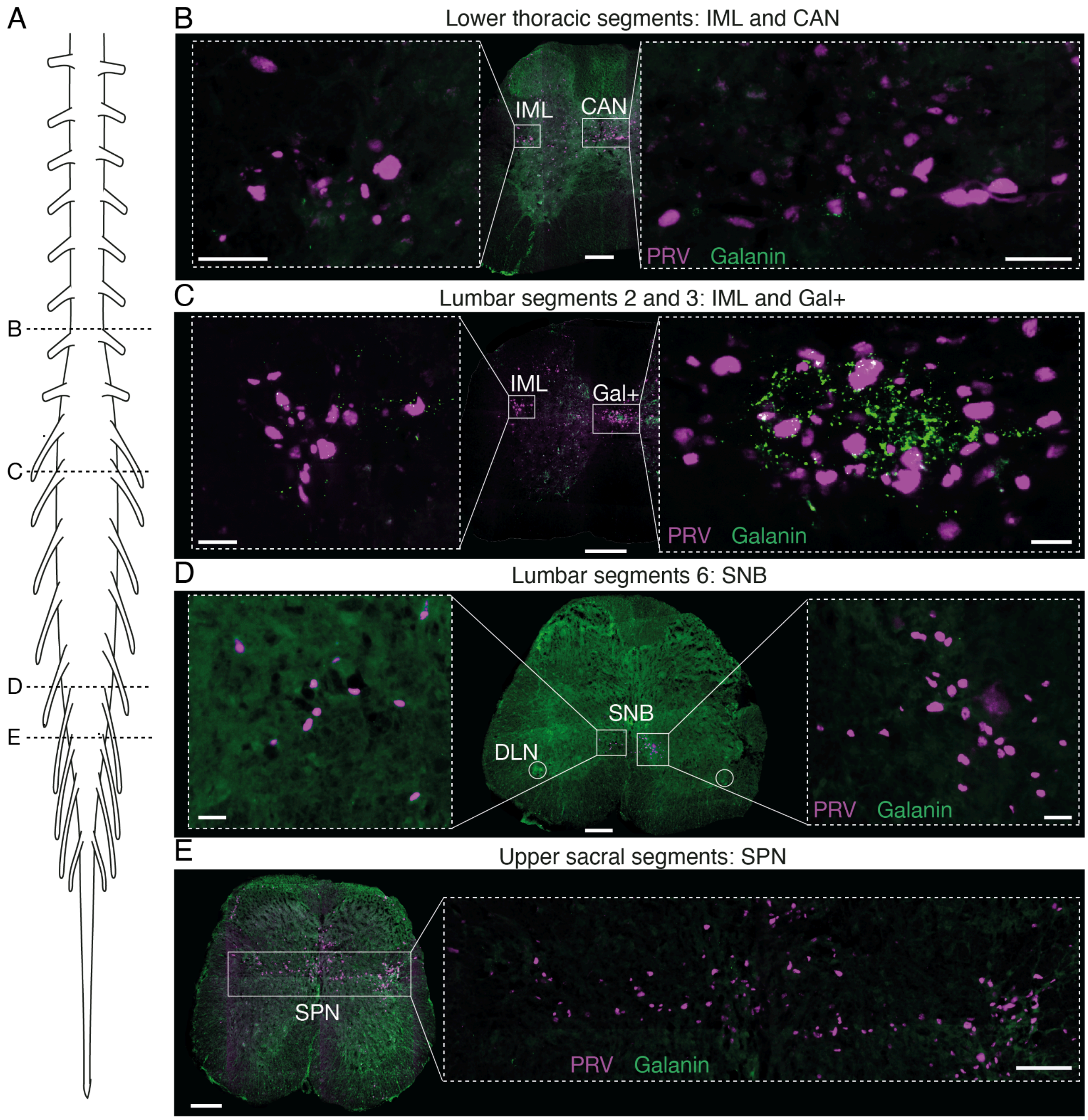


**Supplemental Figure 6: Pseudorabies virus (PRV) injections into the BSM reveal similar anatomical sites as the ones obtained with the injection of AAV-flex-CAG-SynGFP at the location of the Gal+ cells.**

(A) Spinal cord scheme.

(B) PRV-positive labeled cells were found at the location of the intermediolateral (IML) and central autonomic nucleus (CAN) at lower thoracic segments. Scale bar 200um. Scale bar inset 20um.

(C) Prominent PRV labeling was encountered at the central canal and at the IML in L2/3 segments. Scale bar 200um. Scale bar inset 20um.

(D) Clusters of PRV cells were present only in the spinal nucleus bulbocavernosus (SNB), that contains the BSM-MNs, at the L6 segment, but not on the dorsolateral nucleus (DLN, white circle), containing the MNs controlling the ischiocavernosus muscle, which shows the specificity of the PRV approach to trace the MNs innervating the BSM. Scale bar 200um. Scale bar inset 50um.

(E) In the upper sacral segments, PRV labeling could be seen along the spinal parasympathetic centers (SPN). Scale bar 200um. Scale bar inset 100um.

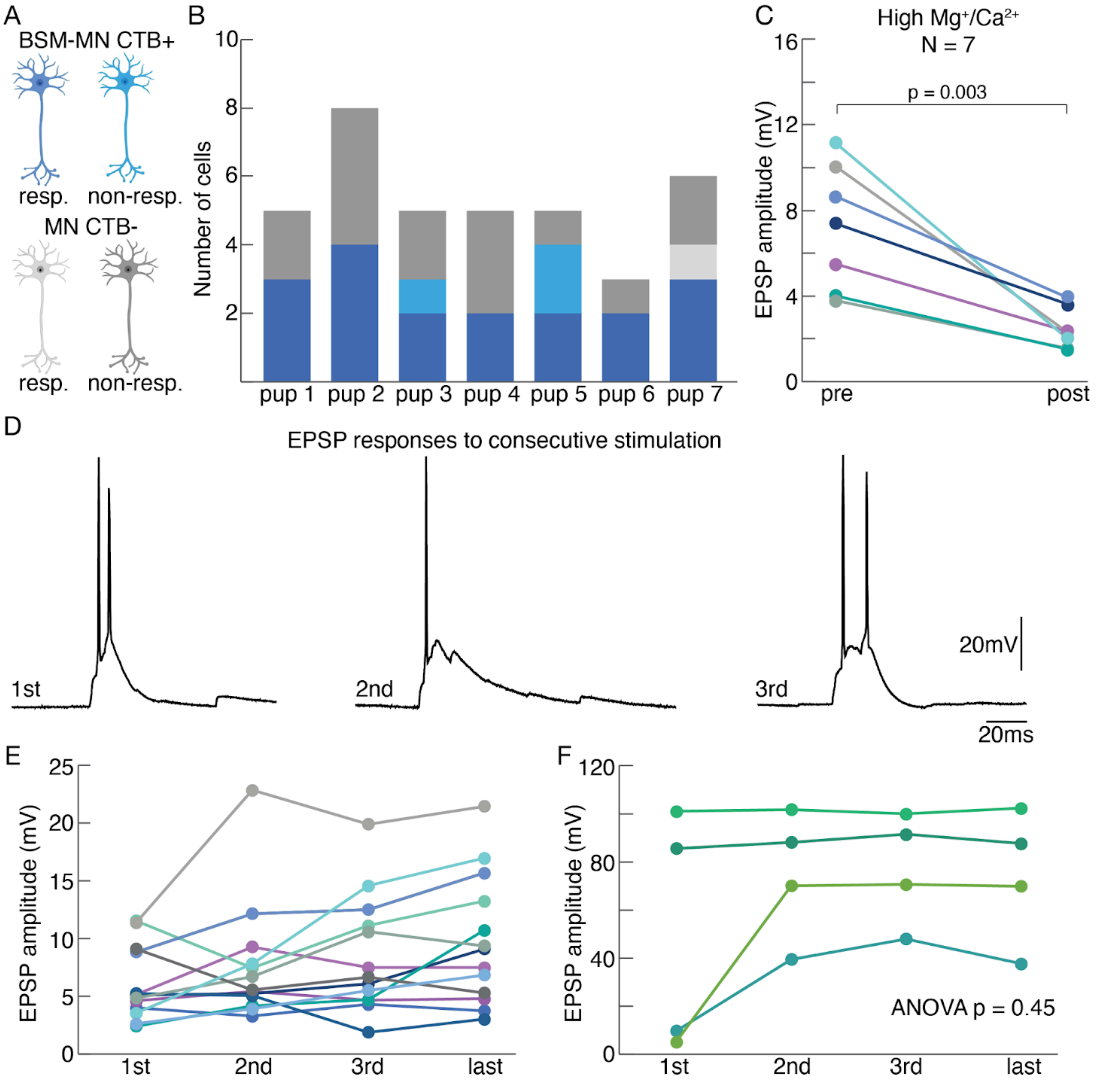


**Supplemental Figure 7: *In vitro* whole cell patching of identified BSM-MNs.**

(A) BSM-MNs were visualized and identified by injecting a CTB-597 into the BSM of Bl6 mice pups. Responses to laser light were compared between CTB+ BSM-MNs (dark blue: responding to laser; light blue: not responding to laser) and CTB- MNs (light gray responding to laser; dark gray non responding to laser).

(B) Distribution of the number of patched MNs per pup depicting responsive (dark blue) and unresponsive (light blue) CTB+ BSM-MNs and responsive (light grey) and unresponsive (dark grey) CTB- MNs.

(C) Light triggered EPSPs amplitudes for CTB+ BSM-MNs before (pre) and after (post) application of High Mg+/Ca2+ application ($p = 0.003$, Student's t-test).

(D) Example recording of a CTB+ BSM-MN depicting light triggered EPSPs upon consecutive laser light application (1st, 2nd, 3rd).

(E) and (F) Light triggered EPSP amplitudes of individual BSM-MNs are plotted for 1st, 2nd, 3rd and last laser light application. Note that consecutive stimulations did not lead to significant different EPSPs amplitudes (ANOVA, $p = 0.45$).

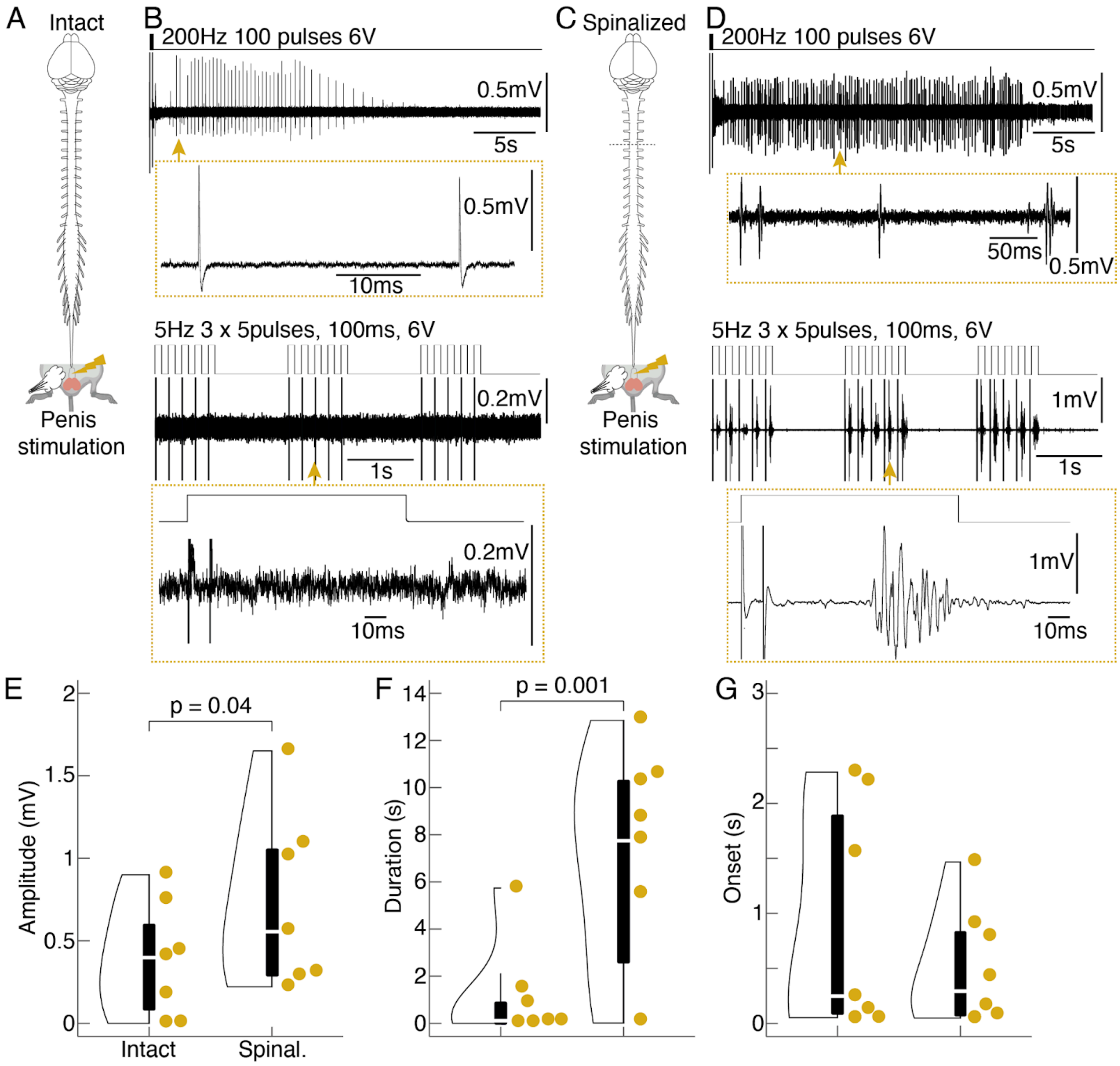


**Supplemental Figure 8: Mechanical and electrical stimulation of the penis leads to more pronounced BSM EMG activity in a spinalized anesthetized *in vivo* preparation**

(A) Experimental design: stimulation of the penis was achieved with either current application using a nerve cuff electrode or with an airpuff, pointed to the penis that was pulled out. The anesthetized preparation was intact, meaning that the connection with the brain was kept.

(B) Example EMG traces obtained in an animal in which the connection to the brain was kept intact and electrical stimulation was applied to the penis. Upper panel: train stimulation. Lower panel: 5Hz stimulation. Note that the train stimulation triggered modest BSM responses while the 5Hz stimulation did not trigger any activity in the BSM EMG.

(C) Same as A but the anesthetized preparation was spinalized, meaning that the connection to the brain was cut.

(D) Same as B but for the spinalized preparation. Note that both the train stimulation and 5Hz stimulation triggered striking BSM responses.

(E) Violin plot comparing the amplitude of BSM responses upon sensory penile stimulation in an intact vs. spinalized (spinal.) preparation (N = 7). Penis stimulation led to significantly bigger BSM amplitudes in spinalized preparations in comparison to the intact preparations. Every dot represents an animal.

(F) Same as E but the duration of BSM responses is plotted. Penis stimulation led to significantly longer BSM responses in spinalized preparations in comparison to the intact preparations.

(G) Same as E but the onset of BSM responses is shown. No significant difference was found.

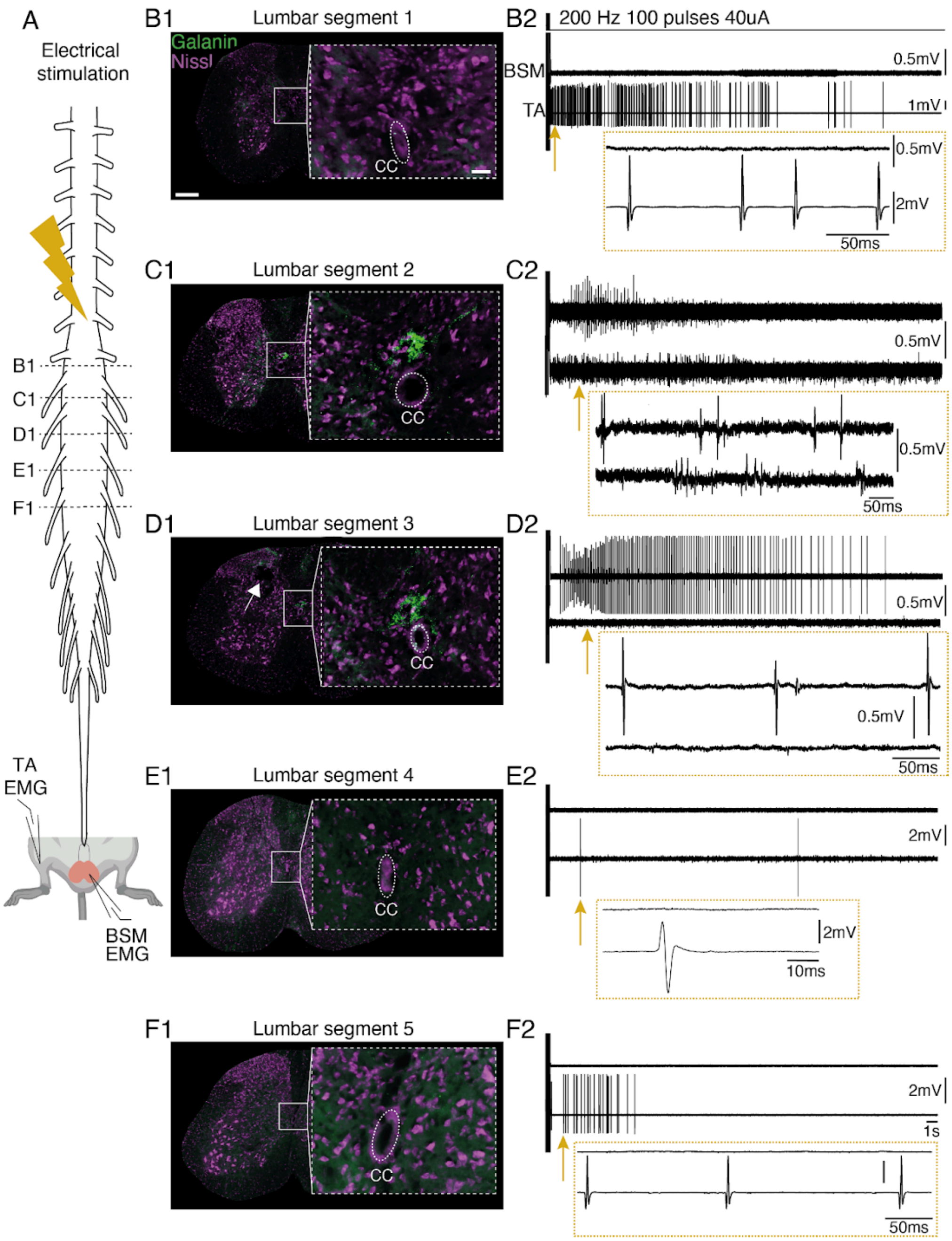


**Supplemental Figure 9: Electrical stimulation along the rostrocaudal lumbar spinal cord in an anesthetized preparation.**

(A) Experimental design: anesthetized animals were clamped into the spinal frame and the spine was opened. A tungsten electrode was inserted along the rostrocaudal axis at different depths and current was applied while BSM activity was monitored in parallel.

(B1 - F1) Representative post-hoc histological sections along the rostrocaudal axis of the spinal cord are shown. Note the electrolytic lesion at the L3 segment (D1). Green: Galanin. Purple: Nissl.

(B2 - F2) Representative electrophysiological traces with BSM (upper trace) and leg (lower trace, TA, *tibialis anterior*) responses triggered at the stimulation site shown in B1 - F1. Note that BSM responses were most prominent at locations (C2, D2) where Galanin labeling was observed post hoc around the central canal (C1, D1) while TA responses randomly occurred along the rostrocaudal axis.

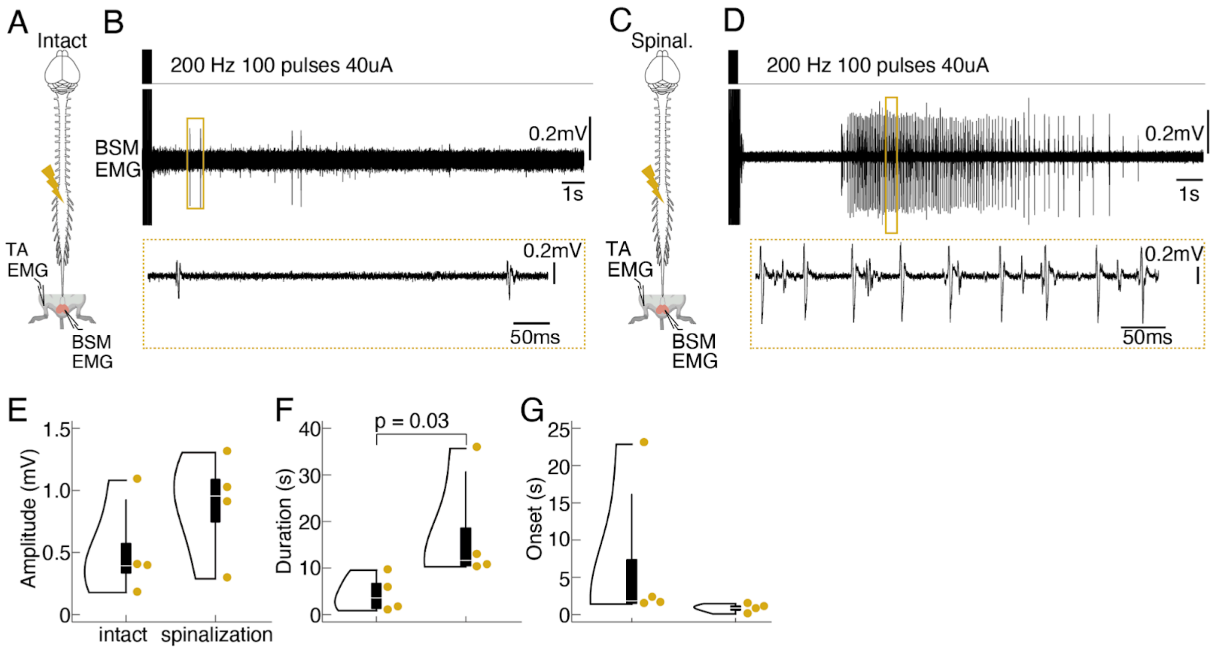


**Supplemental Figure 10: Electrically triggered BSM responses in a spinalized vs non-spinalized preparation.**

(A) Experimental design: electrical stimulations at the location of Gal+ cells in L2-L3 segments (see yellow flash) were conducted in anesthetized mice in which the connection to the brain has been kept intact while in parallel monitoring BSM activity using EMG recordings (N = 4).

(B) Example EMG trace obtained in an animal in which the connection to the brain was kept intact and electrical stimulation was applied with a tungsten electrode.

(C) Same as A, but a spinalization was performed between thoracic segments 5 and 6 (N = 4, same animals).

(D) Same as C, but EMG trace is recorded from an animal in which a spinalization was performed prior to stimulation. Note the difference in BSM activity.

(E) Violin plots showing the BSM EMG amplitude triggered with electrical stimulation in intact anesthetized mice (mean amplitude 0.51 ± 0.19 mv) compared to spinalized mice (0.87 ± 0.21 mV). P = 0.48 resulting from a Mann-Whitney U-Test.

(F) Same as E but the duration of the BSM EMG activity is plotted, which was markedly longer in the spinalized (mean duration 17.33 ± 0.21 s) vs. intact (mean duration 4.41 ± 2.02 s) anesthetized preparations. P = 0.03 resulting from a Mann-Whitney U-Test.

(G) Same as E but the onset of triggered BSM responses is depicted (mean onset for intact 7.0 ± 5.28 s; vs mean onset for spinalized 0.84 ± 0.28 mice). P = 0.05 resulting from a Mann-Whitney U-Test.

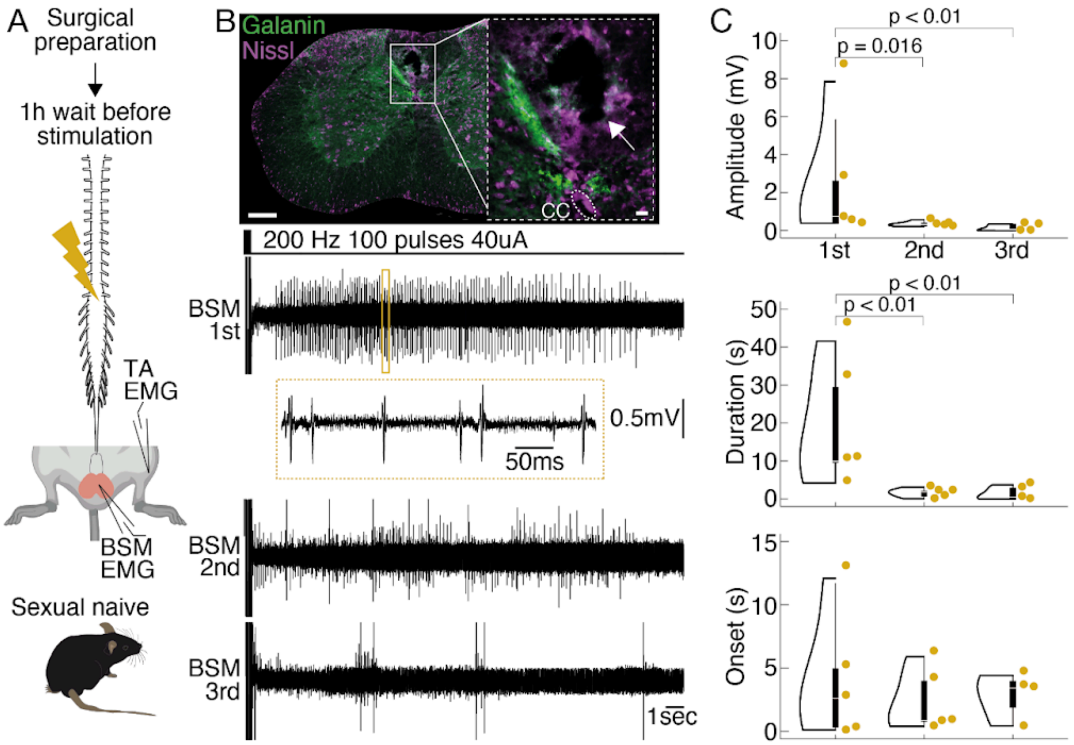


**Supplemental Figure 11: Electrically triggered BSM activity does not depend on the duration of the spinal aperture.**

(A) Experimental design: anesthetized animals (N = 5) were clamped into the spinal frame and the spine was exposed. Before starting with the electrical stimulation protocol, a waiting window of 1h was respected in order to reveal that the observed depression in the BSM activity with repeated trains of stimulation was not due to deterioration of the prep (due to the interval between opening the spinal cord and the application of the last stimulation protocol).

(B) Upper panel: Histological section showing the electrolytic lesion (white flash) that has been placed at the location on which electrical stimulation triggered the highest BSM activity. Green: Galanin. Purple: Nissl. Scale bar 200um. Scale bar inset: 50um. Lower panel: Electrical stimulation protocol (200Hz, 100 pulses, 40uA) led to pronounced BSM activity (BSM 1st) at the location of Galanin clusters. The same depression in BSM activity (as depicted in main Figure 5) was observed during the 2nd and 3rd rounds of stimulation.

(C) Quantitative analysis of BSM responses triggered after 1h of waiting time. Amplitude (upper panel; mean amplitudes: 1st 2.48 ± 1.39 mV; vs. 2nd 0.35 ± 0.06 mV; vs. 3rd 0.13 ± 0.08 mV; 1st vs. 2nd $P = 0.016$ and 1st vs. 3rd $P = 0.0078$ resulting from a Mann-Whitney U-Test) and duration (middle panel; mean durations: 1st 18.9 ± 7.08 s; vs. 2nd 1.61 ± 0.51; vs. 3rd 1.4 ± 0.76 s; 1st vs. 2nd $P = 0.007$ and 1st vs. 3rd $P = 0.008$ resulting from a Mann-Whitney U-Test) of BSM activity decreased upon 2nd and 3rd stimulation rounds while the onset with which BSM activity was elicited remained stable (lower panel; mean onsets: 1st 4.01 ± 2.2 s; vs. 2nd 2.38 ± 1.08 s; vs. 3rd 2.75 ± 0.9 s; $P = 0.98$ resulting from a Kruskal Wallis test).

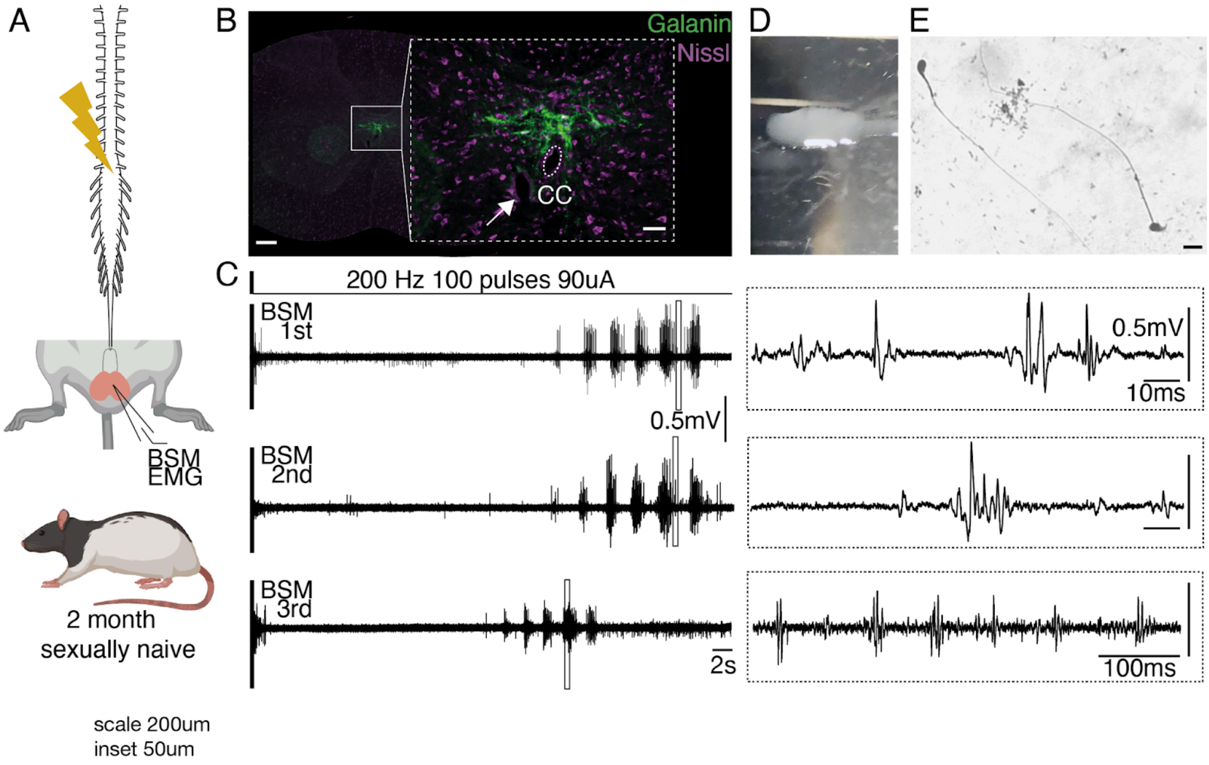


**Supplemental Figure 12: Repeated electrical stimulation of the rat spinal cord at the location of the ejaculation generator leads to stable BSM activity and to the emission and expulsion of sperm.**

(A) Similar to the mice experiment, the tungsten electrode was moved along the rostrocaudal axis of the spinal cord in an anesthetized rat (sexually naive, 2 months) while the BSM activity was monitored in parallel using an EMG.

(B) Histological section of the L3 spinal segment where BSM activity was elicited and ejaculation triggered by current application. Note a similar expression pattern of Galanin (green) around the central canal (CC) when compared to mice. An electrolytic lesion (white arrow) was placed at the shown location. Purple: Nissl stain. Scale bar 200um, inset 50um.

(C) Example BSM EMG trace for the first current application at the location shown in B, note the characteristic rhythmic pattern in the BSM with a long onset. The 2nd and 3rd current application (BSM 2nd, BSM 3rd) led to a very similar activity pattern in the BSM, contrary to mice.

(D) Current applications at the L3/4 spinal segments not only led to a characteristic activity pattern in the BSM, but also to the expulsion of sperm. Cloudy liquid (sperm) was collected on an objective slide which was diluted for post-hoc staining.

(E) Sperm was made visible by performing a Papnicolaou staining protocol. Scale bar 10um.

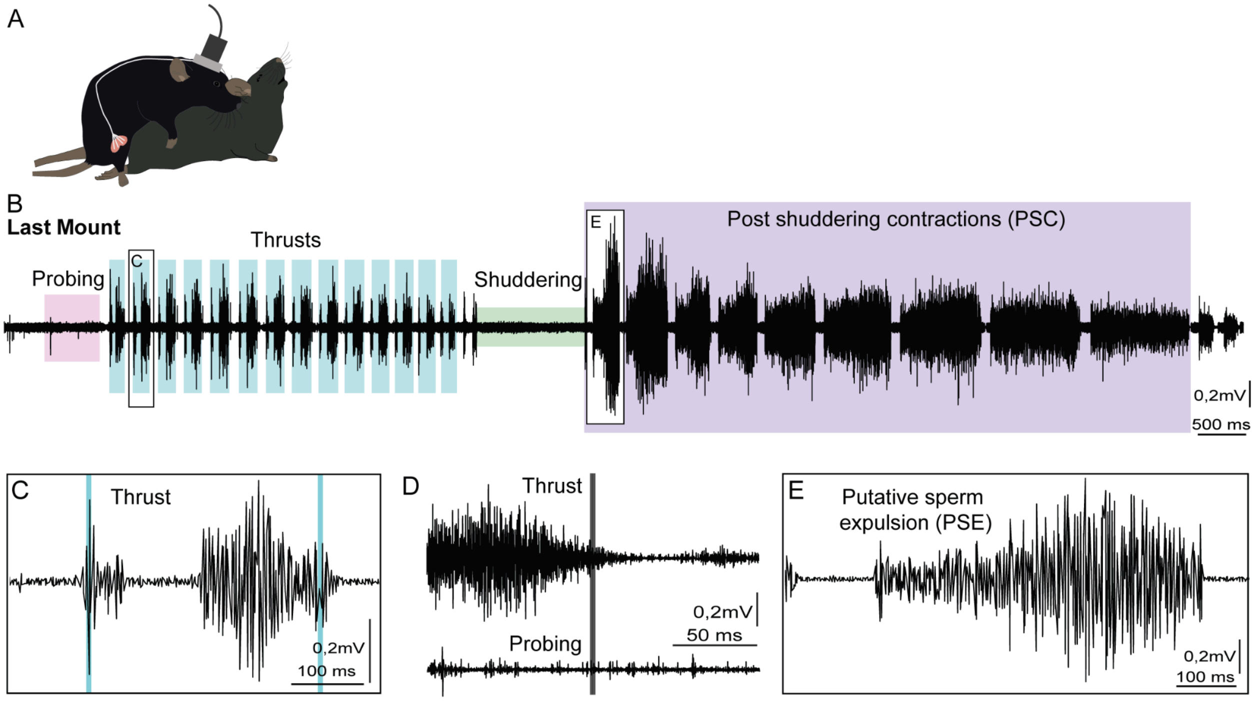


F

| | Amplitude (mV) | Duration PSE (ms) | Duration PSC (s) |
|---|---|---|---|
| Mouse #1 | 0,591 | 270 | 6,83 |
| Mouse #2 | 0,873 | 230 | 6,28 |
| Mean | 0,732 | 250 | 6,55 |

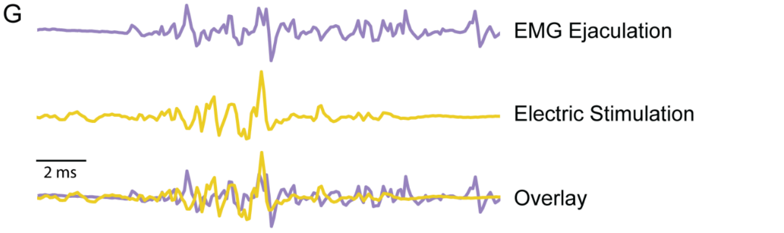


**Supplementary Figure 13 - BSM EMG *in vivo* recordings in sexually behaving animals shows the muscle activity in different phases of the behavior.**

(A) Experimental design: *In vivo* BSM EMG recordings in behaving mice while performing copulatory behavior allowed access to the BSM contraction patterns during the different phases of a sexual encounter (N=2).

(B) EMG trace from the BSM activity in the last mount of a sexual behavior session, showing the muscular activity during Proing, Thrusts, Shuddering and the Post Shuddering Pelvic Contractions.

(C) Close up from the highlighted Thrust in (A), showing the detailed BSM EMG activity. Note the two bursts of activity concordant with the point where the thrust is deeper (first blue trace) and when the thrust is shallower (second blue trace).

(D) Comparison between the thrusts and probing events, aligned to the end of the behavioral bout. Mean EMG trace for all the events in one animal in one recording channel.

(E) Close up from the highlighted Post Shuddering Pelvic Contractions in (A), showing the detailed BSM EMG activity. According to work from McGill and Coughlin, we believe that this is the burst that leads to the expulsion of sperm, hence calling it putative sperm expulsion.

(F) Quantitative analysis of BSM responses in terms of amplitude of the Post Shuddering Pelvic Contractions (mean of 0,732mV), their duration (mean of 6,55s) and the duration of the putative sperm expulsion burst (mean of 250ms) (N=2).

(G) Comparison between the oscillatory pattern of the BSM EMG signal recorded in vivo during expulsion and the BSM EMG signal after electrical stimulation of the Gal+ cells.

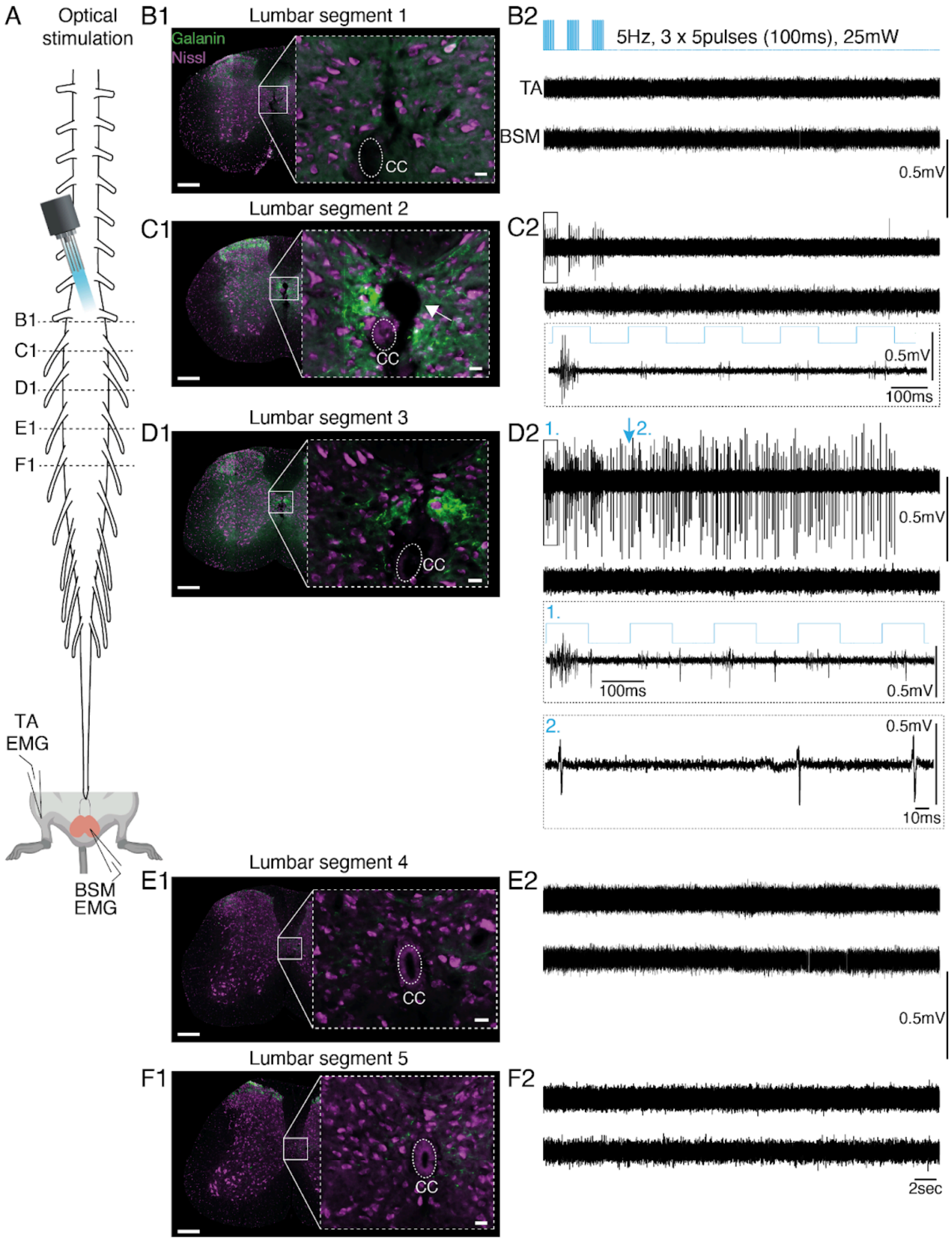


**Supplemental Figure 14: Optogenetic stimulation of the Galanin expressing neurons in an anesthetized preparation.**

(A) Experimental design: anesthetized Gal-ChR2 animals were clamped into the spinal frame, the spine was opened and animals were spinalized. An optical fiber was placed on top of the spinal cord and moved along the rostral caudal axis to deliver blue light while in parallel BSM activity was monitored.

(B1 - F1) Representative post-hoc histological sections along the rostral caudal axis of the spinal cord are shown. Note the electrolytic lesion at the lumbar segment 2 which corresponds to the location where the biggest and longest BSM responses were triggered. Green: Galanin ChR2. Purple: Nissl.

(B2 - F2) Representative electrophysiological traces with BSM and leg (TA, *tibialis anterior*) responses triggered at the optogenetic stimulation site shown in B1 -F1. Note that BSM responses were most prominent at locations (C2, D2) where Galanin ChR2 labeling was observed post hoc around the central canal (C1, D1) while TA responses were never observed.

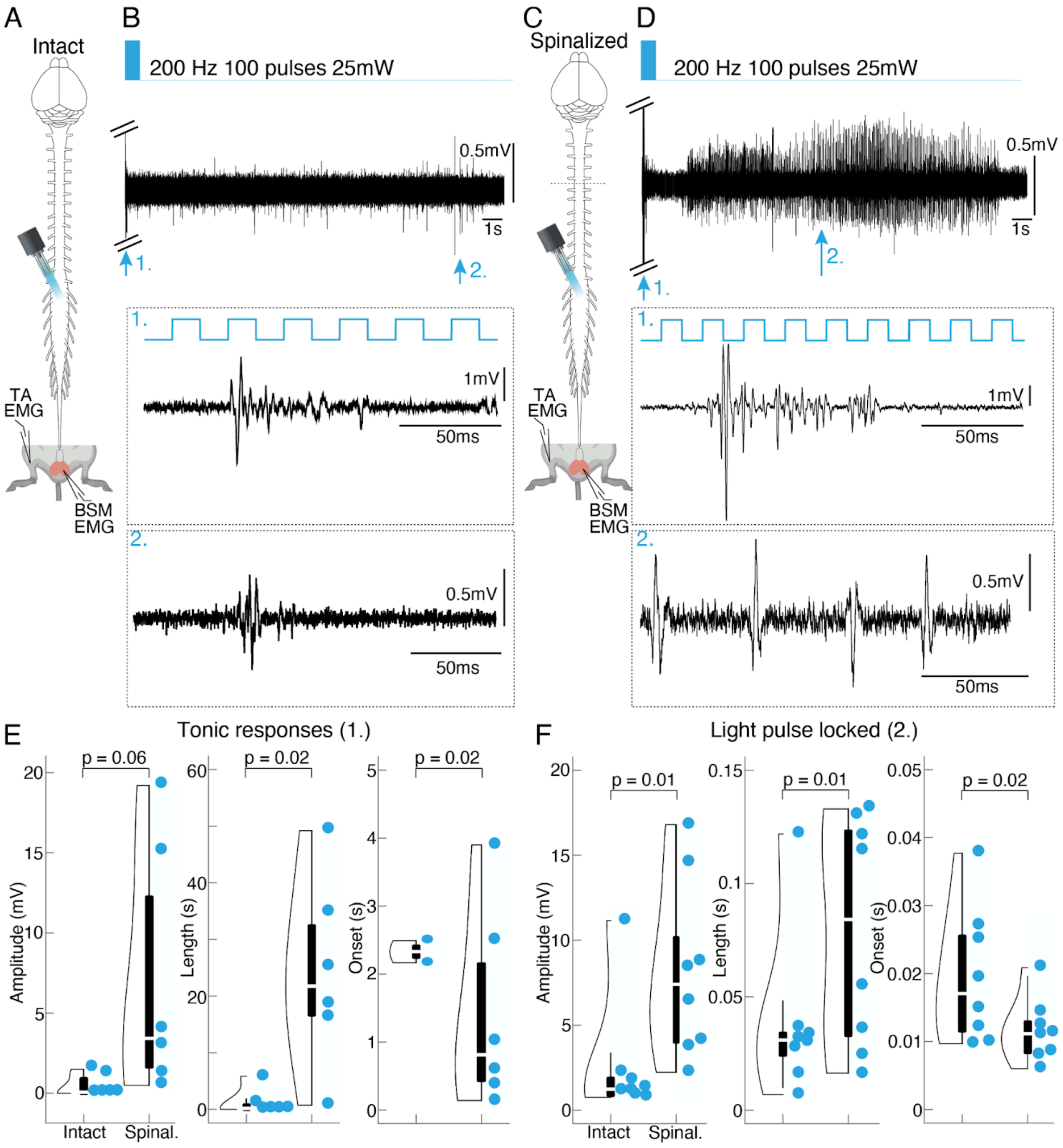


**Supplemental Figure 15: Optogenetic activation of Gal+ cells triggered BSM responses in a spinalized vs non-spinalized preparation.**

(A) Experimental design: optogenetic stimulations were conducted in the Gal+ cells location in anesthetized mice in which the connection to the brain has been kept intact while in parallel monitoring BSM activity using EMG recordings (N = 8; same animals used for intact and spinalization experiments).

(B) Example EMG trace obtained in an animal in which the connection to the brain was kept intact and optogenetic stimulation was applied with an optrode on the spinal cord surface (above the midline).

(C) Same as A but a spinalization was performed between thoracic segments 5 and 6.

(D) Same as B but EMG trace is recorded from an animal in which a spinalization was performed prior to stimulation. Note the difference in BSM activity.

(E) BSM EMG amplitude (left), length (middle) and onset (right) triggered with an optogenetic train stimulation in intact anesthetized mice compared to spinalized mice. Note that amplitudes (mean amplitude intact 0.45 ± 2.6 mV vs. mean amplitude spinalized 7.13 ± 2.6 mV; P = 0.06 resulting from a Student's t-test) and lengths (mean length intact 1.13 ± 6.7 s vs. mean length spinalized 24.1 ± 7.01 s; P = 0.007 resulting from a Student's t-test) were markedly increased after spinalization while the onset with which BSM responses were triggered was shortened after spinalization (mean onset intact 2.32 ± 0.5 s vs. mean onset spinalized 1.42 ± 0.6 s; P = 0.02 resulting from a Student's t-test). Only in 2 out of 8 animals responses were triggered before spinalization.

(F) Same as E but parameters are depicted for the 5Hz optogenetic stimulations (see Figure 5 and supplemental Figure 12). The same phenomena was observed. Light triggered BSM responses showed an increase in amplitude (left; mean amplitude intact 2.5 ± 1.24 mV vs. mean amplitude spinalized 8.11 ± 1.83 mV; P = 0.01 resulting from a Student's t-test) and length (middle; mean length intact 0.04 ± 0.01 s vs. mean length spinalized 0.08 ± 0.02 s; P = 0.05 resulting from a Student's t-test) and a decrease in onset (right; mean onset intact 0.02 ± 0.003 s vs. mean onset spinalized 0.01 ± 0.003 s; P = 0.02 resulting from a Student's t-test) after the connection to the brain was cut.

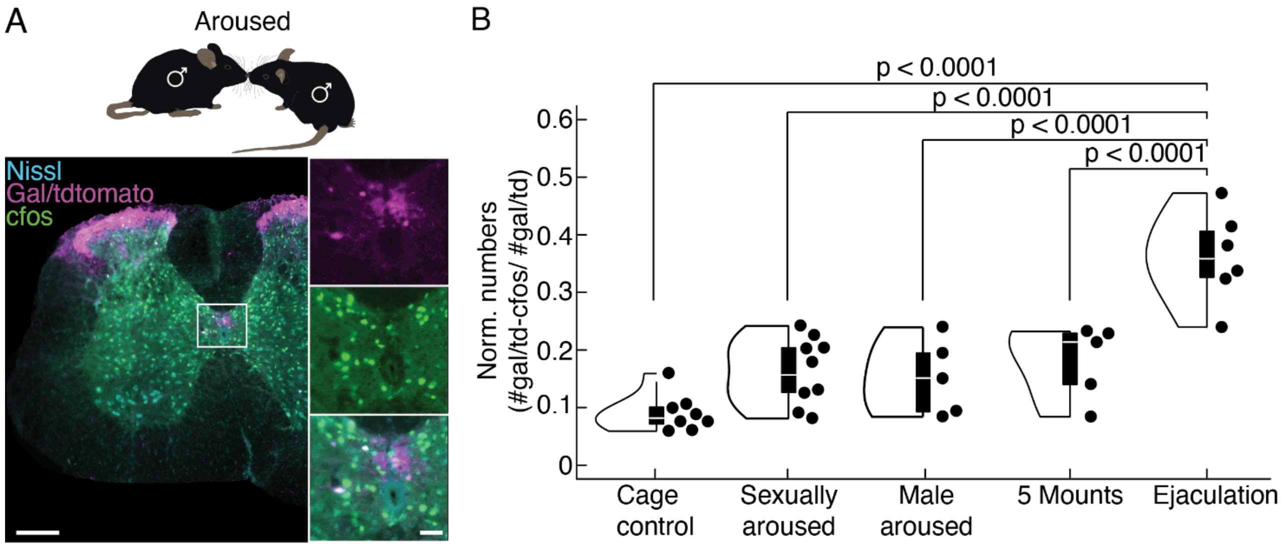


**Supplemental Figure 16: Activation of the Gal+ population with different social interactions, namely a male-to-male interaction.**

(A) Upper panel: Males in the Male aroused group were allowed to interact with a naive young male (5 weeks old) for 10min or 10 attack bouts (N = 5). Lower panel: Example spinal cord section at the L2/3 segments.

(B) Normalized cell numbers (number of double labeled cells, cfos+ and Gal+; divided by the total number of Gal+ cells) for the five groups (groups depicted in Figure 6 with the addition of the Male aroused group). Depicted p values from Student's t-test (violin plots elements: see Methods).

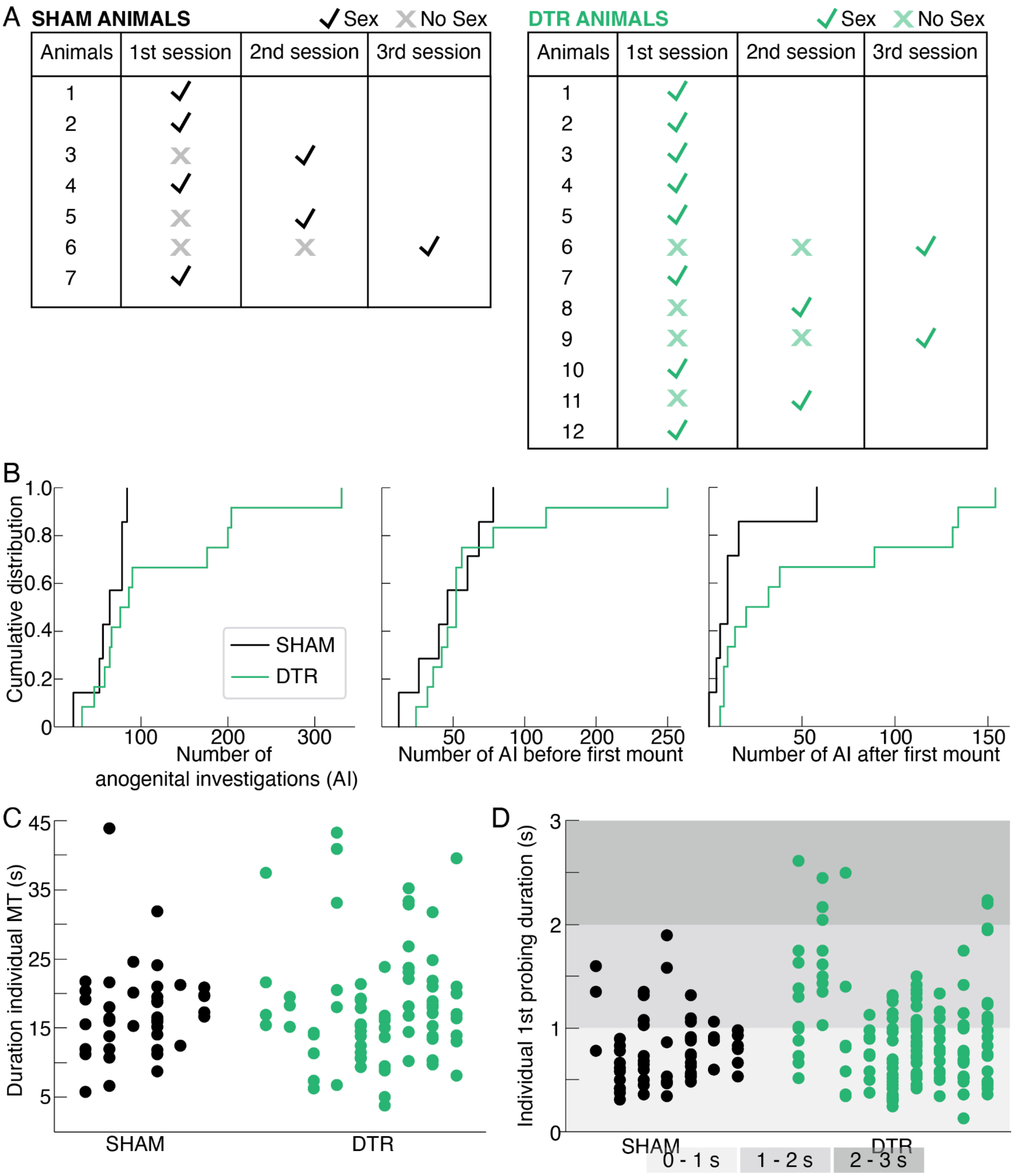


**Supplemental Figure 17: SHAM and DTR animals have similar behavior patterns over several sessions.**

(A) Animals were tested for 3 weeks in a row to assess their capability to perform sexual behavior after surgery. Both SHAM (left) and DTR (right) animals took different amounts of sessions to initiate the behavior, but eventually all showed sexual behaviors (mounts, probing, thrustings, and in most cases ejaculation) in the span of three weeks.

(B) Cumulative distribution relative to the number of anogenital investigations shown by SHAM and DTR animals. Left panel all anogenital investigations are plotted, middle panel number of anogenital investigations before the first mount and right panel number of anogenital investigations after the first mount are depicted. No significant difference was found using the Mann-Whitney-Wilcoxon two-sided test ($p=0.17$, $p=0.6$ and $p=0.07$, from left to right respectively) or the two-sample Kolmogorov-Smirnov test ($p=0.15$, $p=0.96$ and $p=0.26$).

(CB) Duration of individual mounts with thrusts (MT, in seconds) per animal. SHAM and DTR animals show identical values.

(C) Duration of the first probing event per mount (MP, in seconds) per animal. DTR animals show a higher amount of mounts with longer probing durations (namely with more than 2 seconds duration).

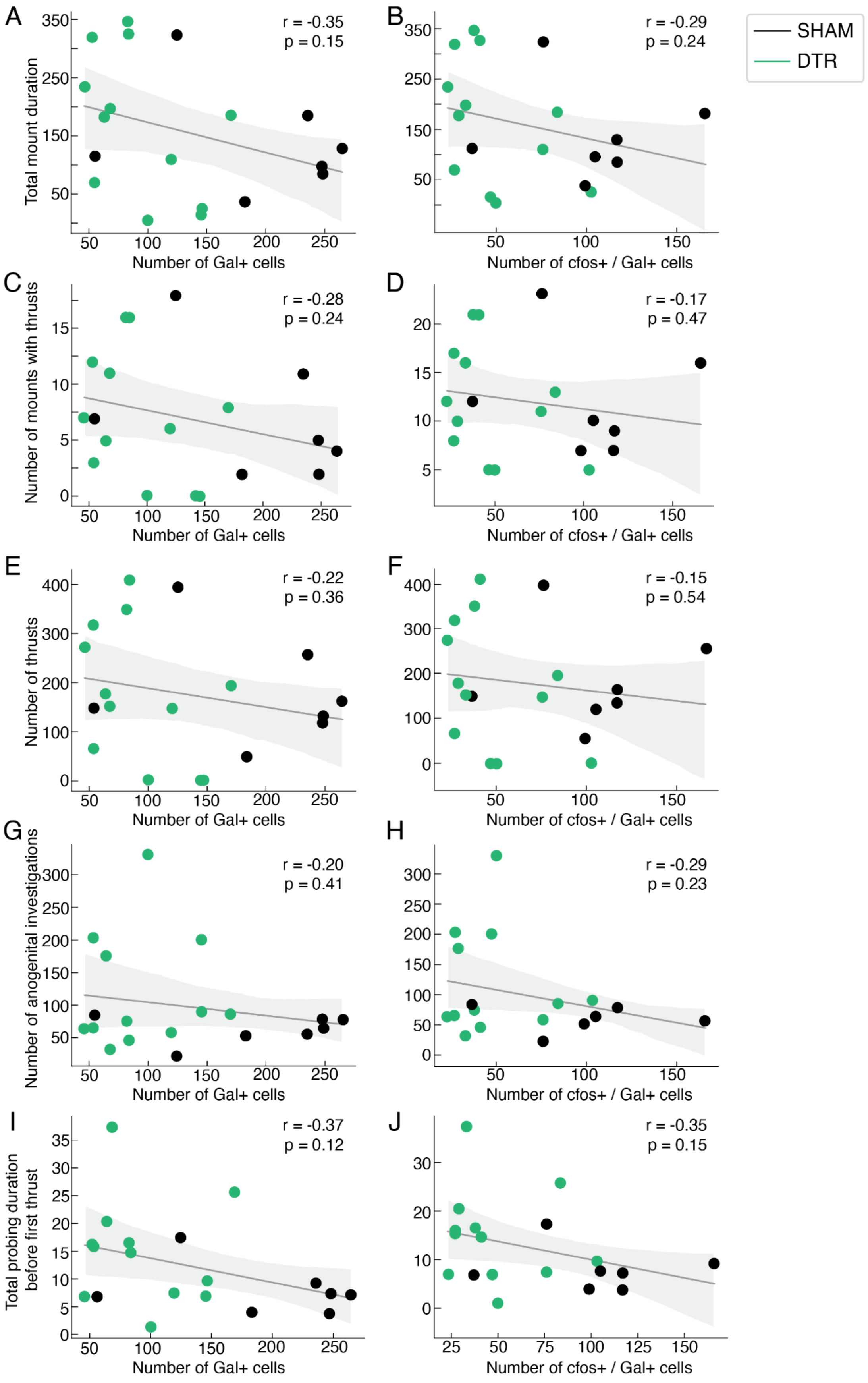
A
r = -0.35
p = 0.15
Total mount duration
Number of Gal+ cells
B
r = -0.29
p = 0.24
Number of cfos+ / Gal+ cells
SHAM
DTR
C
r = -0.28
p = 0.24
Number of mounts with thrusts
Number of Gal+ cells
D
r = -0.17
p = 0.47
Number of cfos+ / Gal+ cells
E
r = -0.22
p = 0.36
Number of thrusts
Number of Gal+ cells
F
r = -0.15
p = 0.54
Number of cfos+ / Gal+ cells
G
r = -0.20
p = 0.41
Number of anogenital investigations
Number of Gal+ cells
H
r = -0.29
p = 0.23
Number of cfos+ / Gal+ cells
I
r = -0.37
p = 0.12
Total probing duration before first thrust
Number of Gal+ cells
J
r = -0.35
p = 0.15
Number of cfos+ / Gal+ cells

**Supplemental Figure 18: No correlation between sexual behavior measures and amount of Gal+ cell ablation.**

(A) Total mount duration is plotted against the number of Gal+ cells in SHAM (black) and DTR animals (green). Every dot represents an individual animal. Note that there is no significant correlation between the Gal+ cell numbers and the total mount duration.

(B) Total mount duration is plotted against the number of Gal+ and cFos+ cells in SHAM (black) and DTR animals (green). Every dot represents an individual animal. There is no correlation between the number of double positive cells (cFos+ and Gal+) and the total mount duration.

(C) Same as A but for the number of mounts with thrusts.

(D) Same as B but for the number of mounts with thrusts.

(E) Same as A but for the number of thrusts.

(F) Same as B but for the number of thrusts.

(G) Same as A but for the number of anogenital investigations.

(H) Same as B but for the number of anogenital investigations.

(I) Same as A but for the total probing duration before the first thrust.

(J) Same as B but for the total probing duration before the first thrust.

### Supplemental Movies

#### Supplemental Movie S1

Light evoked BSM-MN activity leads to characteristic pelvic floor movements

#### Supplemental Movie S2

Pelvic floor movements during ejaculation in a sexually behaving male mouse

#### Supplemental Movie S3

Conservation of erection behavior in a DTR animal.

#### Supplemental Movie S4

Erection behavior in a SHAM animal.
